# An Empiricist’s Guide to Modern Coexistence Theory for Competitive Communities

**DOI:** 10.1101/640557

**Authors:** Casey M. Godwin, Feng-Hsun Chang, Bradley Cardinale

## Abstract

While most ecological theories have historically invoked niche differences as the primary mechanism allowing species coexistence, we now know that species coexistence in competitive communities actually depends on the balance of two opposing forces: niche differences (ND) that determine how species limit their own growth rate versus that of their competitor, and relative fitness differences (RFD) that establish competitive hierarchies among species. Several different empirical methods have been proposed for measuring ND and RFD in order to make predictions about coexistence of species, yet it remains unclear which method(s) are appropriate for a given empirical study and whether or not those methods actually yield the same information. Here we summarize and compare five different empirical methods, with the aim of providing a practical guide for empiricists who want to predict coexistence among species. These include two phenomenological methods that estimate ND and RFD based on observing competitive interactions among species; two mechanistic methods that estimate ND and RFD based solely on information about species’ resource requirements; and a fifth method that does not yield ND and RFD but describes the impacts of those forces within communities. Based on the specific requirements, limitations, and assumptions of each approach, we offer a series of decision steps that can be used to determine which method(s) are best for a given study system. In particular, we show there are important tradeoffs between mechanistic methods, which require detailed understanding of species niches and physiology but are more tractable experimentally, and phenomenological methods which do not require this detailed information but can be impractical for some study designs. Importantly, we show that although each method can be used to estimate ND and RFD, the methods do not always yield the same values. Therefore we caution against future syntheses that compile these estimates from different empirical studies. Finally, we highlight several areas where modern coexistence theory could benefit from additional empirical work.

## Introduction

Throughout most of the history of community ecology, it has been assumed that niche differentiation among species is primary biological mechanism that can offset the negative impacts of interspecific competition on species coexistence (Volterra 1931, Gause 1934, Tilman 1982, Leibold 1995, Chase and Leibold 2003). This idea originated when Volterra (Volterra 1931) introduced a dynamic model of competition that became the foundation for the competitive exclusion principle, which states that if two species have identical niche requirements then one of them will inevitably become locally extinct (Gause 1934). The competitive exclusion principle led to two conclusions about coexistence in competitive communities: 1) species will coexist only if they are limited by different resources (or consumers) at the same location and time, or if they partition resources (or consumers) in space or time and, as a result, 2) ecosystems should contain only as many species as there are limiting resources (or consumers) (Rescigno and Richardson 1965, MacArthur and Levins 1967, MacArthur 1970, Tilman 1977, Leibold 1995). Nearly all subsequent hypotheses to explain coexistence have argued that biodiversity exists because of such niche differences among species.

While niche partitioning remained the theoretical basis for understanding species coexistence, empiricists and theorists struggled to reconcile the rich biodiversity that exists in many of the world’s ecosystems with the prediction that competitive communities should contain only as many species as there are limiting resources (or consumers) (Hutchinson 1961, Oksanen et al. 1981, Huisman and Weissing 1999). But starting in 2000, theories of species coexistence began to undergo a major revision. In 2001, Hubbell published The Unified Neutral Theory of Biodiversity (Hubbell 2001), which argued that patterns of biodiversity in nature can be explained by a simple model that does not invoke niche differences among species. According to Hubbell’s theory, species coexist not because they are different, but because their demographic parameters are identical, or nearly so, such that the consequences of their interactions are ‘neutral’ (i.e. essentially equal among all species). Based on this neutral theory, Hubbell argued that the biodiversity we observe in nature can be explained by a series of stochastic events that cause some populations to become dominant while others exhibit random walks toward extinction.

Even as Hubbell was developing his neutral theory, Chesson (2000) was completing a ground-breaking theory of coexistence that would provide a framework for integrating the niche and neutral perspectives on biodiversity. Chesson’s coexistence framework was built on his insight into the invasibility criterion: a pair of species can coexist only if each species is capable of invading a steady-state population of its competitor. Chesson showed how a species’ long-term growth rate when invading a resident species can be decomposed into two general terms, which he called stabilizing and equalizing forces. Stabilizing forces cause species to limit their own growth rate more than they limit the growth rate of other species (intra > interspecific competition). These stabilizing forces, also known as niche differences (ND), occur when species partition limiting resources in space or time, or when they experience differential consumption by consumers. In contrast, equalizing forces minimize differences in competitive abilities among species. Equalizing forces, which have also been called relative fitness differences (RFD), are the result of inherent variation in biological traits such as minimum requirements for shared resources or consumers, differential resistance to consumers, or differences in potential growth rates (Adler et al. 2007, Levine and HilleRisLambers 2009, HilleRisLambers et al. 2012).

Chesson showed it is the balance of these two forces – RFDs that establish competitive hierarchies, and NDs that prevent competitive exclusion – that ultimately determines whether species maintain non-negative long-term growth rates in competitive communities (Chesson 2000). For a pair of species to coexist, ND must be sufficiently large to offset and stabilize the competitive hierarchies generated by RFDs. When two species exhibit identical niches (ND equals 0), their RFD alone determines the competitive hierarchy and which species will become extinct. It has subsequently been shown that Hubbell’s neutral theory represents a specific, limiting case of Chesson’s coexistence theory where NDs and RFDs are both zero, causing the outcome of competition to be approximated by a random walk toward extinction (Adler et al. 2007). Stabilizing and equalizing forces have been identified in both fluctuation-dependent mechanisms (e.g. storage effects) and fluctuation independent mechanisms of coexistence (e.g. competition for a limiting resource) (Miller and Klausmeier 2017, Barabas et al. 2018). Thus, Chesson’s inequality provides a general framework for predicting species coexistence.

Since the development of Chesson’s theory, much attention in ecology has turned towards the empirical estimation of ND and RFD in order to determine how these two forces contribute to coexistence in real communities. As a growing number of empiricists have tried to quantify ND and RFD in their individual study system, the number of different empirical approaches proposed for doing so has also grown. However, the various methods for implementing modern coexistence theory in empirical studies have been derived from different models of species interactions, make different assumptions, and use different experimental designs. Therefore, it remains unclear which method(s) are best suited for a given study, whether the methods give comparable estimates of ND and RFD, and whether the methods actually give the same prediction regarding coexistence. If Chesson’s theory is to become widely implemented in empirical studies and in applied contexts, we need a ‘users guide’ to help ecologists determine which empirical approach meets their needs.

Here we provide a summary and comparison of four methods that have been proposed to measure ND and RFD empirically, and a fifth method that does not give estimates of ND and RFD but has been used to predict coexistence based on Chesson’s theory. In Part 1 of our paper, we describe the theoretical background of each method, illustrate how it can be implemented empirically, and ask whether the methods yield the same estimates of ND and RFD and make the same predictions regarding coexistence. In Part 2 of the paper we provide a list of decision steps to guide empiricists in selecting the most appropriate method(s) for their study system and contrast the methods in terms of the amount of effort required to implement them. In Part 3, we discuss the main advantages and disadvantages of the methods and make some suggestions for future empirical work on coexistence theory.

## Part 1. Summary of Five Empirical Methods for Implementing Chesson’s Theory

In this part of the paper we first review the fundamentals of Chesson’s theory that are most relevant to empirical work on competition and coexistence. We then summarize each of five empirical methods that have been used to measure niche differences (ND) and relative fitness differences (RFD) by (i) explaining how the method relates to Chesson’s theory, (ii) showing how the method can be implemented empirically, and (iii) highlighting the method’s key limitations and assumptions. At the end of Part 1, we use numerical simulations to compare the five methods to determine if they give the same estimates of ND and RFD, and to assess whether they yield the same prediction for coexistence based on the invasibility criterion.

### 1.1 Brief review of Chesson’s theory

When Chesson first introduced his theory for coexistence, he did not prescribe a specific empirical approach or experiment that should be used to estimate ND and RFD in real biological communities. Instead, he used a phenomenological model of competition to show how the mutual invasibility criterion, a common prerequisite for coexistence, depends on how each species limits their own growth rate versus that of their competitor (Chesson 1990). Specifically, he showed that the criterion for mutual invasibility can be expressed as an inequality involving both ND and RFD (Equation 1).

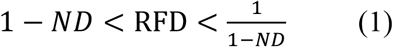

In this inequality, the term 1-ND represents the degree of niche overlap (*ρ* in Chesson (1990)), which ranges from zero when species do not share any resources to one when the resource requirements of species are identical. RFD represents the ratio of competition-free fitness among the two species (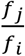 in Chesson (1990)). If this equality is not satisfied, then one of the species is unable to maintain long-term, positive growth rates and will go locally extinct.

Because ND and RFD are not terms that cannot be quantified directly from experiments or observations, Chesson showed how these forces can be derived from the classic Lotka-Volterra competition model. In this model, the *per capita* growth rate of species *i* is a function of both intraspecific and interspecific competition as described by Equation 2:

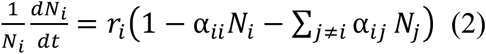

In Equation 2, *N_i_*is the density of species *i,* and *r_i_* is the intrinsic per capita growth rate of species *i*. The intra-specific competition coefficient *α_ii_* describes the *per capita* effect of species *i* on the *per capita* relative growth rate of species *i* and is equal to the inverse of the carrying capacity (*K_i_*) for species *i*. The inter-specific competition coefficient *α_ij_*describes the *per capita* effect of species *j* on the *per capita* relative growth rate of species *i*. Equations 3 and 4 relate the inter- and intra-specific interaction coefficients from the Lotka-Volterra model to ND and RFD:

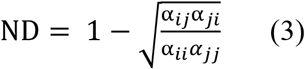

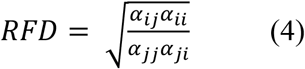

ND and RFD from Equations (3) and (4) can then be used in Equation 1 to predict coexistence. Because Chesson derived ND, RFD and the inequality for mutual invasibility based on the Lotka-Volterra competition model, we use the same approach to explain four of the empirical approaches described below and show that a fifth approach is ultimately not compatible with Chesson’s derivation.

### 1.2 Method based on the parameterized Lotka-Volterra model

#### 1.2.1 Theoretical background

Since Chesson originally used the Lotka-Volterra model to explain his criterion for coexistence, the most obvious empirical approach for estimating ND and RFD is to parameterize the Lotka-Volterra competition model (Equation 2) using data collected from experiments or time-series observations from natural ecosystems. The inter- and intra-specific interaction coefficients (*α_ii_ and α_ij_*) from the Lotka-Volterra competition model can be used to estimate ND and RFD using Equations 3-4.

#### 1.2.2 Empirical approaches

Empirical implementation of the Lotka-Volterra model requires estimating six different parameters used in Equation 2: intrinsic per capita growth rate of each species (*r_i_*and *r_j_*), per capita intra-specific competition coefficients (α_*ii*_ and α_*jj*_), and per capita inter-specific competition coefficients (α_*ij*_ and α_*ji*_). The simplest way to parameterize the Lotka-Volterra model from experiments would be to use three treatments for each pair of species: a time-series of each species grown alone as a monoculture and one time-series representing a co-culture of the two species (Figure 1). From each monoculture time series, the empiricist needs to measure the population density of each species over time, from low density to steady-state. From these time series, the empiricist can estimate the maximum per capita growth rate of each species (*r_i_*), which occurs as the species’ density approaches zero, and the steady-state population size of each species in monoculture (*K_i_*).

**Figure 1.**
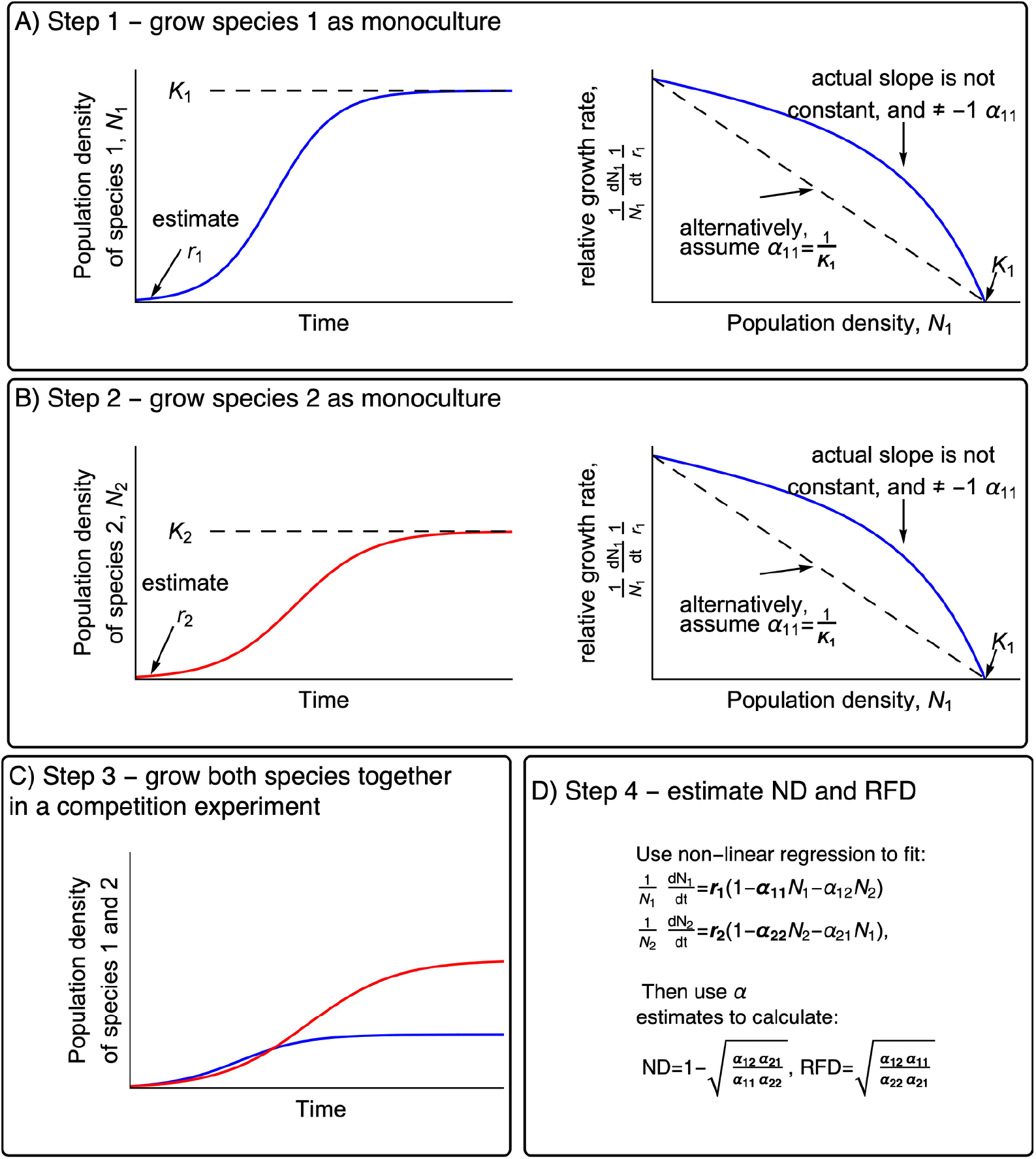
Conceptual plots illustrating how to use the Lotka-Volterra method to estimate ND and RFD for use in Chesson’s inequality (Equation 1). In each panel, unknown parameters are displayed in regular typeface and previously-estimated parameters are listed in bold typeface. In panels a and b, the left-hand plots show the time course of the experiment. In panels a and b the right-hand plots show the relative growth rate as a function of population density – the slope of this relationship is equal to the intraspecific competition coefficient (sign reversed).

Empiricists have two options for using the monocultures to estimate the intraspecific competition coefficients (*α_ii_*). The first option is to estimate these interaction coefficients using the slope of each species’ relative growth rate (scaled on its maximum growth rate *r_i_*) versus its population density (Figure 1A and 1B, right). This slope has the opposite sign of the intraspecific interaction coefficient *α_ii_*. However, in practice this slope is unlikely to be fixed across all population densities. As a result, intra- and inter-specific competition coefficients measured in competition experiments are likely to vary with population densities (Abrams 1980). An alternative strategy is to use the assumption that intraspecific competition coefficients (*α_ii_*) are equal to 1/*K_i_*, which comes directly from the Lotka-Volterra model in Equation 2. This approach yields estimates of *α_ii_* that cannot, say, be used in Equation 2 to re-create transient dynamics observed in time series, but can be used in Equations 3 and 4 to estimate ND and RFD and predict coexistence using Chesson’s inequality (Equation 1).

Next the empiricist would perform a competition experiment in which both of the species are introduced to habitat at low density and the population density of each species is measured over time (Figure 1C). From these time series, the empiricist must use non-linear regression to parameterize the interspecific interaction coefficients (α_*ij*_ and α_*ji*_) by substituting the parameter estimates from the monocultures into Equation 2. Finally, the empiricist can use all four interaction coefficients to compute ND and RFD using Equations 3 and 4.

#### 1.2.3 Limitations

A key assumption of this approach is that the intra- and inter-specific competition coefficients are fixed with respect to population sizes of either species. In other words, the first individual and the last individual added to a population have the same per capita effect on the growth rates of its own species or that of its competitor. This assumption is not always met in real biological communities where intra- and inter-specific competition coefficients can depend on species’ densities (Smith-Gill and Gill 1978). The assumption also does not apply when the mechanisms leading to competition are driven by non-linear dependence on resources. Examples include functional responses of consumers to prey density and non-linear dependence of growth rates on abiotic resource availability (e.g. the Monod function). Figure A1 in the Supporting Information shows that, when applied to numerical simulations based on a well-known consumer resource model parameterized with real biological data, intraspecific coefficients measured in monoculture near equilibrium lead to inaccurate predictions regarding coexistence. However, when the intraspecific interaction terms are replaced by 1*/K_i_*the method yields accurate predictions. Therefore, in those situations where competition coefficients are fixed with respect to population size, or can be measured at low population sizes of each species, then the empirical approaches can be used to estimate ND and RFD.

### 1.3 Sensitivity method

The second method for estimating ND and RFD, the sensitivity method, is similar to the Lotka-Volterra method in that it is phenomenological and requires information from direct competition experiments.

#### 1.3.1 Theoretical background

The sensitivity method quantifies the proportional reduction in a species’ growth rate when invading a steady-state population of its competitor (Carroll et al. 2011, Narwani et al. 2013). In this method, the maximum growth rate of each species in monoculture (*µ_i_*) and when invading a steady-state population of the competitor species (*µ_ij_*) are used to calculate each species sensitivity to interspecific competition (*S_i_*) using Equation 5:

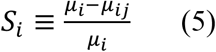

Carrol and others (2011) and others have shown that ND is proportional to the geometric mean of these sensitivity measures, whereas RFD represent variation around the mean:.

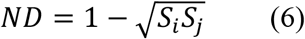

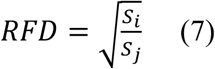

A species’ sensitivity to competition is jointly determined by ND and RFD (Carroll et al. 2011, Narwani et al. 2013). Specifically, greater ND between the two species reduces the impact of interspecific competition so that *S_i_*will approach zero. Greater RFD, on the other hand, causes species to be asymmetrically affected by competition such that one species’ sensitivity increases while the other’s decreases. While Carroll and others (2011) verbally argued that this method is compatible with Chesson’s theory, in Appendix A we show explicitly how this method relates to Chesson’s theory and prove that it is identical to Equations 3 and 4 when interspecific interaction coefficients of the Lotka-Volterra model are evaluated only at the steady-state population density of the resident species.

#### 1.3.2 Empirical approaches

The sensitivity method uses a combination of monocultures and pairwise invasion experiments to quantify the reduction in each species’ growth rate caused by a steady-state population density of the other species at its carrying capacity (Figure 2). The experiment by Narwani et al. (2013) provides an example for how to implement the sensitivity method empirically. Their experimental system involved species of freshwater green algae growing under controlled conditions in the laboratory. They grew each species as a monoculture in fresh growth medium, starting from low biomass and allowing the populations to reach their carrying capacity. From these time series, they quantified the per capita maximum growth rate of each species as a monoculture, which occurs when the focal species is at low density (*μ_i_* and *μ_i_*). After each species reaches its carrying capacity, they introduced the other species from low density (e.g. 0.01% of *K*) and quantified the per capita growth rate of each species when invading the other (*μ_ij_* and *μ_ji_*). Finally, for each pair of species, they used these growth rates to calculate the sensitivity metrics (*S_i_* and *S_j_*) using Equation 5 and used those sensitivity metrics to calculate ND and RFD using Equations 6 and 7.

**Figure 2.**
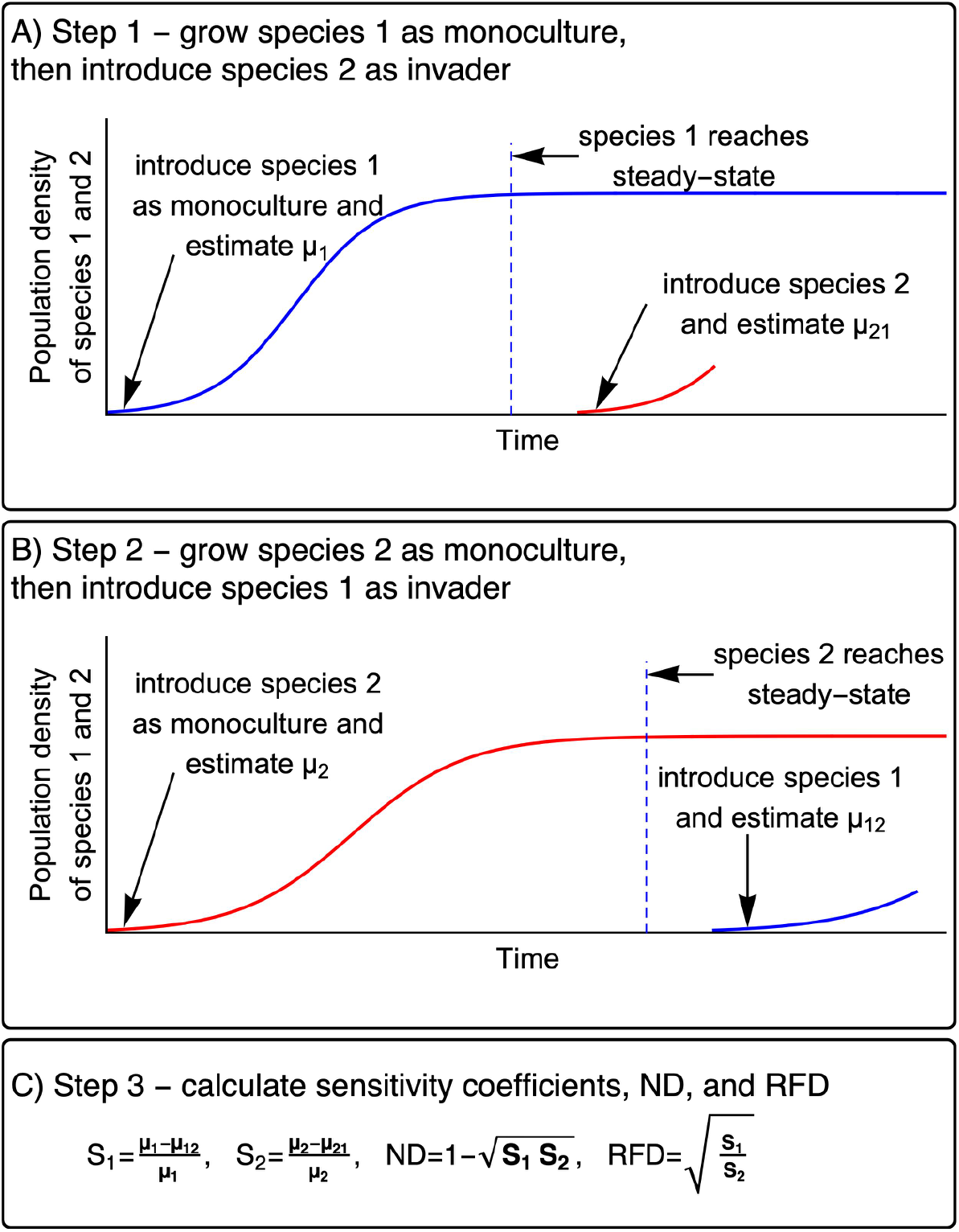
Conceptual plot depicting how to implement the sensitivity method in an experiment.

#### 1.3.3 Limitations

Using the sensitivity method requires one to perform mutual invasibility experiments, which are only practical for organisms whose population growth rates can be measured over tractable periods of time. Mutual invasibility experiments are harder to apply to organisms that grow slowly (e.g. trees) since it would take a long time to acquire time series of species densities needed to estimate per capita growth rates. Also, it is important to note that the invasion growth rates (*μ_ij_* and *μ_ji_*) must be measured when the invader population density approaches zero. Under this condition, intra-specific competition is negligible, and the resident species’ density is near steady-state. If the growth rate of the invader species were measured at greater density of the invader species or lower density of the resident species (i.e. long after invasion), then the *S_i_*would be affected by both intra- and inter specific competition. The resulting predicting regarding species coexistence would be incorrect.

### 1.4 Parameterizing MacArthur’s consumer resource model

The third method to estimate ND and RFD from empirical data is to parameterize MacArthur’s consumer-resource model (MacArthur 1970) then use these parameters to calculate ND and RFD using Chesson’s original derivation (Chesson 1990, 2000). This method is different from both the Lotka-Volterra and sensitivity methods because it does not rely on experiments where the species are grown together in order to quantify how the species influence each other’s growth rates. Instead, this method works by parameterizing a mechanistic model that describes species interactions, then reorganizing those parameters to estimate ND and RFD for assessing Chesson’s inequality.

#### 1.4.1 Theoretical background

MacArthur’s consumer resource model describes how species consume and thus compete for two or more prey resources (MacArthur 1970). The model is composed of differential equations representing the growth of each consumer species as a function of resource densities (Equation 8) and a differential equation (or set) that describes the population dynamics of each prey resource and their mortality due to consumption by the consumers (Equation 9).

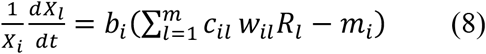

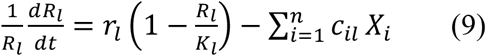

In this model X_4_is population density of the consumer species Y and @_M_is population density of the prey resource Z. The term *b_i_* represents the effect of prey consumption on the growth rate of the consumer, *r_l_* is the maximum per-capita growth rate of prey resource *l*, *K_l_* is the carrying capacity for the prey species *l*, *w_il_* represents the increase in consumer population density for each unit of prey resource *l* consumed. The term c*_il_* is the resource capture rate by consumer *i* on resource *l* and *m_i_* is the density-independent mortality for consumer species *i*. Chesson showed that, by implementing a time-scale separation technique, parameters in MacArthur’s consumer resource model can be used to calculate ND and RFD using Equations 10 and 11 (Chesson 1990, 2000):

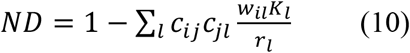

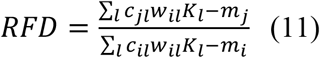

The estimates of ND and RFD from this method can then be used to evaluate Chesson’s inequality and predict coexistence (Equation 1).

#### 1.4.2 Empirical approaches

Using MacArthur’s consumer resource model to estimate ND and RFD for a pair of species requires quantifying 1) the per capita consumption rate of each consumer species on each prey resource (*c_il_*, units of prey consumed per unit prey density); 2) the per capita maximum growth rate and carrying capacity of each prey resource when no consumers are present (*r_l_* and *K_l_*); and 3) the yield of consumer population density or biomass relative to each unity of prey consumed (*w_il_*). Because we are not aware of any empirical studies that have parameterized the MacArthur model for the purpose of estimating ND and RFD, we describe the experimental approach that would be required (Figure 3). First, the empiricist would need to identify or define the prey resources that are available to the consumer species. Each prey resource would be inoculated or planted at low density into an environment free of other prey resources and consumers, then the population density would be measured over time in order to estimate the per capita maximum growth rate of the prey (*r_l_*, which occurs as the prey population density approaches zero) and its carrying capacity (*K_l_*, which occurs when the prey growth rate approaches zero). Next, the experimentalist would need to introduce each consumer species into several different densities of each prey resource growing as a monoculture. Under those different prey resource densities, the experimentalist would measure the per capita consumption rate of prey resource by the consumer species (*c_il_*) and the yield of consumer density or biomass per unit prey resource consumed (*w_il_*).

**Figure 3.**
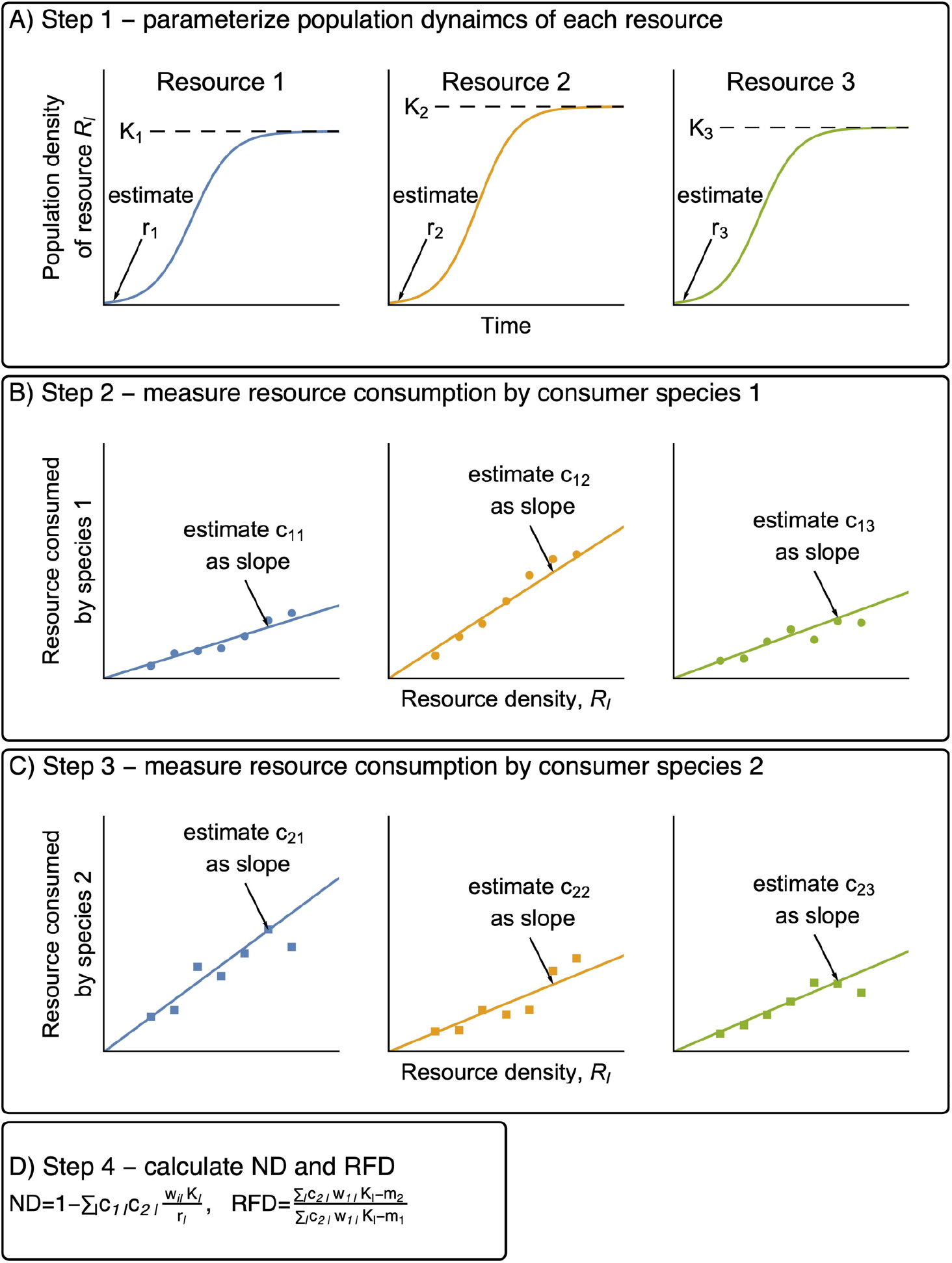
Conceptual plots depicting how the method based on Mac Arthur’s CRM could be implemented. The yield term (*w_il_*, increase in consumer units per unit prey resource consumed), can be estimated by measuring these changes for each combination of consumer and resource.

The precise number of parameters to be estimated depends on the number of prey resources considered by the model. For example, for two consumer species and three prey resources (Figure 3), the hypothetical experiment requires 18 parameters to be quantified: 3 different maximum per capita growth rates and 3 carrying capacities of the prey resources (*r_l_* and *K_l_*, *l* = 1 to 3), 6 per capita consumption rates (*c_il_*; *i*= 1 and 2, *l*= 1 to 3), and 6 yields (*w_il_*; *i*=1 and 2, *l*=1 to 3). These parameters can then be used in Equations 10 and 11 to obtain ND and RFD, which can subsequently be used in Equation (1) to predict coexistence.

#### 1.4.3 Limitations

The MacArthur’s consumer resource model gives a more mechanistic understanding of competitive interactions among species, and allows one to predict coexistence for pairs of species without needing to grow them together in a competition experiment. However, these desirable properties come with greater number of experimental treatments compared to the Lotka-Volterra and sensitivity methods. In particular, this method requires as many consumption experiments as there are resources, and each of these experiments involves measuring consumption rates at a range of resource species densities (Figure 3 B and C). While this constraint does not impact the ability of the method to predict coexistence under defined conditions, it could limit the extent to which those predictions can be applied to natural environments where the number of potential prey species is large.

### 1.5 Parameterizing Tilman’s consumer resource model

Like the method based on MacArthur’s consumer-resource model, the method based on Tilman’s consumer resource model does not require species to be grown together in a competition experiment (Letten et al. 2017). However, unlike the method based on MacArthur’s CRM, the method based on Tilman’s CRM is specific to abiotic resources that are controlled by a constant rate of supply and do not have their own intrinsic growth rate (i.e. a chemostat).

#### 1.5.1 Theoretical background

Letten and others (2017) showed how a consumer resource model (Tilman 1977) can be reorganized to a Lotka-Volterra form in order to estimate ND and RFD and assess mutual invasibility using Chesson’s inequality. This method is based on Tilman’s two-species consumer resource model for two essential and non-substitutable resources (Tilman 1977). In this model, one set of differential equations describes the growth of each consumer species as a function of resource availability (Equation 12) and another set of equations describes the dynamics of abiotic resources and their depletion due to uptake by the consumer and dilution (Equation 13).

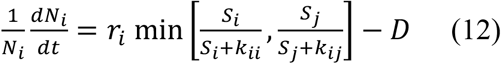

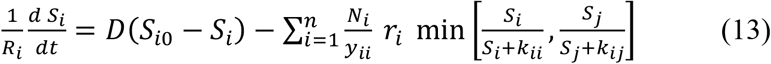

In this model, *N_i_* is the population density of species *i*, *r_i_* is the maximum per capita growth rate of species *i*, *y_ii_* is the yield of species *i* on resource *i*, and *k_ii_* is the half saturation constant for growth of species *i* on resource *i*. The term *S_i0_* is the external supply concentration for resource *i*, *S_i_* is the concentration of resource *i* in the environment, and *D* is equal to both the supply rate of resources and the density-independent loss rate for both species.

Letten et al. (2017) showed how the parameters from Tilman’s CRM can be used to calculate ND and RFD. First, the empiricist must determine which species is limited by each resource (e.g., using Resource-Ratio theory (Tilman 1982)). This requires comparing the supply ratio for the two resources against the R*s for each species at the pre-determined dilution rate. As such, the following equations are specific to only the condition where species 1 is limited by resource 2 and species 2 is limited by resource 1.

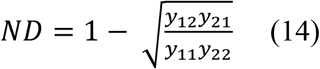

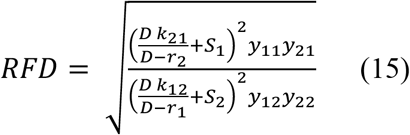

Equations 14 and 15 can be used to estimate ND and RFD and then used in Chesson’s inequality (Equation 1) to predict species coexistence.

#### 1.5.2 Empirical approaches

To illustrate how this method could be implemented empirically, we describe the approach that Tilman first used to parameterize his model (Tilman 1977). In the chemostat environment associated with Tilman’s model, the abiotic resources or nutrients are delivered to the experimental system at a constant supply rate that matches the density-independent mortality rate (Tilman 1977, 1982). First, he measured the growth kinetics for two algae species when limited by two different essential resources (silicate and phosphate). He inoculated each species as a monoculture into growth medium containing a range of concentrations of the limiting resource (either silicate or phosphate) with all other resource in excess. From these time series of population densities, he quantified the growth rate of each species as a function of the concentration of the limiting resource following the Monod model (Figure 4). For each species, this yields estimates of half saturation constants (*k_ij_*) for each resource and a single maximum per capita growth rate for both resources (*r_i_*). Next, Tilman quantified the yields (*y_ij_*) of each species on each resource by measuring the elemental content of a known number of cells. By following the same approach, an empiricist can quantify the 10 parameters needed to define Tilman’s model.

**Figure 4.**
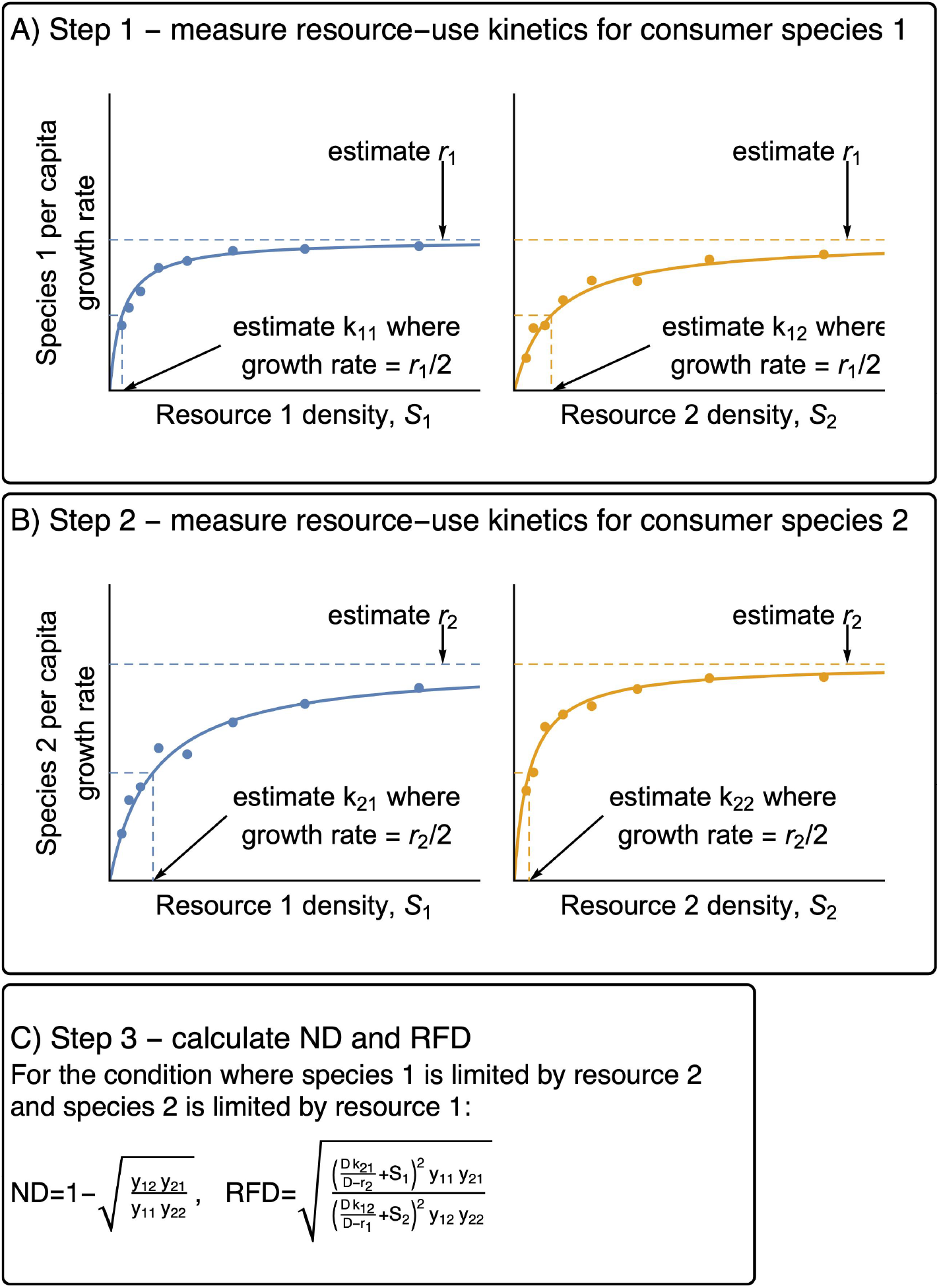
Conceptual plot depicting how to parameterize the method based on Tilman’s consumer resource model. Panels A and B show the experiments needed to parameterize the maximum growth rates and Monod half-saturation constants for growth on each resource, separately for each species. The yield of each species on reach resource (*y_ii_*) can be estimated by measuring the amount of resource consumed by a known number of individuals.

As shown by Letten et al. (2017), the parameters described above can be used to predict coexistence under different resource supply ratios and dilution rates in a chemostat. However, the form of the equation used to get ND and RFD depends upon the empiricist knowing whether or not the supply ratio is outside the ratio of 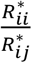 and 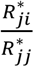 for the two species, as determined by resource-ratio theory (Tilman 1982).

#### 1.5.3 Limitations

The method using Tilman’s consumer resource model requires an empiricist to know precisely which resources the species compete for, which limits its applicability to many real scenarios and ecosystems where the identity of limiting resources and the supply rates may not be known. Additionally, the resource supply concentrations must be fixed and the supply rates must be equal to the density-independent loss rate, which can only be achieved in certain experimental settings like chemostats. Another important note is that the elemental content of organisms like algae is not constant and can vary as a function of growth rate, nutrient availability, and other factors (Sterner and Elser 2002). Non-constant yields due to luxury uptake and variable internal stores (Grover 1991) have important consequences for competition, but it has yet to be determined how this plasticity affects estimation of ND and RFD.

### 1.6 Negative frequency dependence (NFD) method

The final method that we summarize, the negative frequency dependence method (NFD), has not been proposed as a means of obtaining estimates of ND and RFD that are directly compatible with Chesson’s inequality (Equation 1). In fact, in Supporting Information C, we explicitly show that the NFD method cannot be used to derive estimates of ND and RFD that are consistent with Chesson’s theory. However, the NFD method can be used to predict coexistence using the criterion of mutual invasibility. Moreover, the NFD method has been proposed as a way to interpret stabilizing and equalizing forces from Chesson’s original theory, and the method has been used to illustrate the impacts of ND and RFD in manipulative experiments (Adler et al. 2007, Levine and HilleRisLambers 2009).

#### 1.6.1 Theoretical background

The NFD method quantifies the change in the per capita growth rate of a species as a function of its frequency in a community (Adler et al. 2007, Levine and HilleRisLambers 2009). Here the frequency of a species refers to the proportion of total biomass or individuals in a community belonging to that species. This method makes the key assumption that the community is saturated with respect to total species densities, thus a frequency of 1 represents a steady-state monoculture at its carrying capacity. At all other frequencies, the community composition need not be at a steady-state, but it is assumed that any increase in the population density of one species will be offset by a decrease in population density of another species. Using this assumption, the slope of the NFD relationship has been used to reflect the difference between intra-versus inter-specific competition (Adler et al. 2007). Increasing species *i*’s frequency means that individuals of species *i* will compete more with individuals of its own kind than with individuals of other species, and will thus experience stronger intraspecific competition than interspecific competition. Therefore, if intra-specific competition is greater than inter-specific competition, the species affects its own growth rate more than it affects the growth rate of other species, and the NFD slope should be negative.

The NFD method is most often used as a graphical approach for understanding the balance of ND and RFD (Figure 5). According to Adler et al. (2007), a more negative NFD slope represents a stronger stabilizing force, which they argue is proportional to the ND in Chesson’s inequality. Similarly, they argue that the difference between species’ growth rate in the absence of stabilizing forces is the equalizing force, proportional to RFD. In Supporting Information C we show that this NFD slope is not equivalent to ND and that the difference in elevation between species’ NFD relationships is not RFD. Although the NFD approach does not yield estimates of ND and RFD that are consistent with Equation 1, this method can still be used to make predictions about coexistence based on Chesson’s mutual invasibility criterion. Specifically, if each species has a positive growth rate at a frequency approaching 0 then the mutual invasibility criterion is satisfied, and coexistence is assured. Adler and others (2007) showed how both the slope and elevation of the NFD plot are needed to accurately determine whether this condition is met.

**Figure 5.**
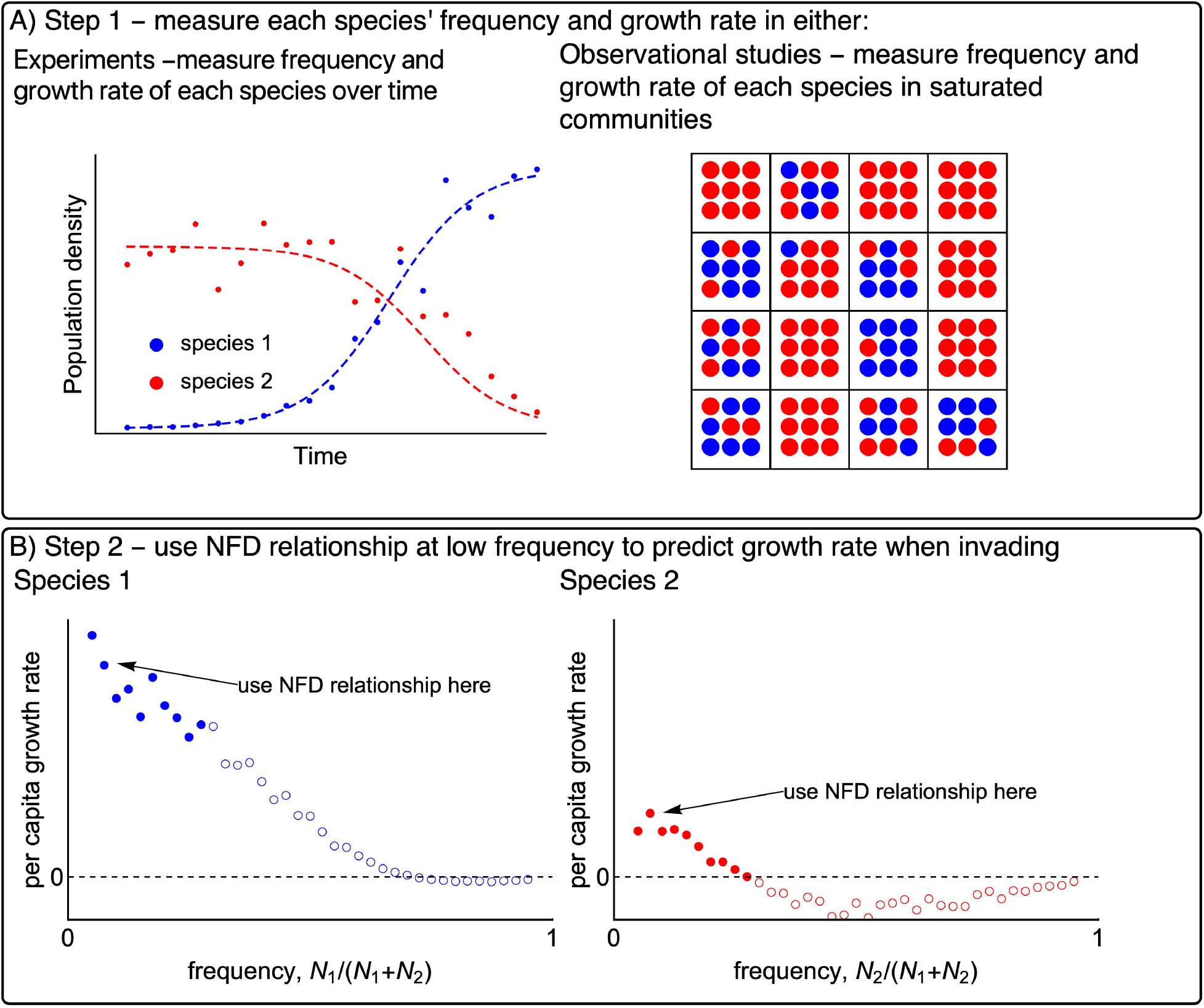
Conceptual diagram depicting how the NFD method could be implemented, either through and experiment or using observational data from different communities. Panel A shows two options for obtaining pairs of each species’ frequency and its growth rate in a saturated community. The first option is to track population densities over time in one or more competition experiments. The Second option is to obtain pairs of frequency and growth rate from different Communities or habitat patches in a natural ecosystem. Panel B depicts how the data from either experiments or observational studies would be used to estimate the growth rate when frequency approaches zero.

#### 1.6.2 Empirical approaches

The NFD method could be implemented using either experiments or observations from natural ecosystems (Figure 5A). Using the experimental approach, an empiricist could growth both species together and measure their densities over time. First, the empiricist would need to ensure that total community density or biomass was saturated. One way to do this would be to use invasion experiments in which the resident species is grown to steady-state and then the invader species is introduced at low density. Using the invasion approach guarantees that the community is saturated since any decrease in the resident’s population density is met with an increase in the density of the invader. Next, the empiricist could use the time series for each species’ density to calculate the per capita growth rate and frequency of each species at each time point. Alternatively, the NFD method could be implemented using observational data from natural ecosystems (Adler et al. 2010). This approach could allow an empiricist to estimate frequency dependence for species that are not easily manipulated (e.g. trees). To use this approach, an empiricist would quantify the per capita growth rate and the relative frequency of the species in different habitat patches or along ecological gradients. Although this approach has not been applied empirically to make predictions regarding coexistence (but see Yenni et al. 2017), it is one of only two methods reviewed here that do not require manipulative experiments.

Having obtained pairs of growth rate and frequency from either experiments or observational studies, the empiricist can construct plots of the NFD relationship (Figure 5B). This relationship can be used to estimate the growth rate when each species approaches frequency of zero. If either of the two species does not have a positive growth rate when rare, then the pair will not coexist. Since the NFD method does not yield estimates of ND and RFD that can be used in Equation 1, the utility of this method is completely dependent on its ability to accurately predict mutual invasibility (i.e. growth rates at frequency of 0). In many cases, this prediction will be based on observations made at species frequencies greater than zero. As long as the relationship between a species’ frequency and its growth rate is linear, the NFD slope and elevation can theoretically be used to predict whether both species will have positive growth rates when rare, thus meeting the mutual invasibility criterion. If the NFD relationship is not linear, then the NFD method can give inaccurate predictions (Figure 5B, see Limitations below). Indeed, even when applied to numerical simulations from the simple Lotka Volterra model, the NFD slope is not constant across species frequencies in a saturated community (Supporting Information Figure C1).

Levine et al. (2009) demonstrated how the NFD method can be implemented experimentally. In their study with 10 species of grassland plants, they manipulated the relative frequency of each focal species by varying the proportion of seeds belonging to the focal species versus all other species. At the end of the growing season, they quantified the growth rate of each species in their study plots by multiplying the number of seeds belonging to each species by the proportion of those seeds that were viable the next year. They then quantified the slope of NFD by plotting the growth rate of each species against its frequency in the initial community. Although the slope of NFD is not equal to Chesson’s ND and the difference in intercepts is not equal to RFD, the authors showed that the effect of niche differences on growth rates can be removed by experimentally maintaining each species’ density at a constant, non-equilibrium level that is not subject to competition from other species. Their experiment showed that effectively removing niche differences among species (even without measuring them) led to dominance by the species with the highest per capita growth rates. In other words, in the absence of ND the outcome of competition was determined by RFD. It is important to note that while this approach is based on Chesson’s inequality, it does not require measuring ND and RFD. Similarly, other studies have measured the slope of NFD as evidence for the importance of stabilizing forces, but did not directly interpret the slope as ND or the intercepts as RFD (Yenni et al. 2017).

#### 1.6.3 Limitations

Despite some of the desirable aspects of the NFD method in terms of empirical approaches (above), it has three key limitations. First, unlike the other four methods summarized in this paper, the NFD method does not yield estimates of ND and RFD. This may not be a concern if the purpose of the study is simply to predict species coexistence and does not focus on explaining why certain pairs coexist while other pairs do not. Second, the NFD method assumes that the community density is saturated across the range of species’ frequencies observed. Meeting this assumption in experiments requires sufficiently long time series to show that total biomass of a community is fixed. In observational studies based on natural ecosystems, it might not be possible to ensure that total biomass is saturated.

The third limitation of the NFD method is that the relationship between a species’ frequency and growth rate is often non-linear (Figure 5). In Appendix C, we show the NFD method can lead to incorrect predictions about species coexistence when applied to systems with non-linear relationships between species’ growth rates and densities. If the slope and elevation of the NFD plot are evaluated over a narrow range of species frequencies, and those data were used to extrapolate to predict growth rate as frequency approaches zero, then the method could make inaccurate predictions about mutual invasibility and coexistence. If the relationship between each species’ frequency and its growth rate is not linear, then an empiricist must adequately describe the relationship to account for the non-linearity. This means that for an empiricist to use the NFD method, they would need to either 1) measure the growth rate of each species across the full range of frequencies to establish that the growth rate of each species is linearly related to its frequency or 2) evaluate the growth rate of each species when rare (i.e. directly demonstrate mutual invasibility). Both of these options would dramatically increase the effort required but may be necessary in systems where only observational studies are possible.

### 1.7 Do the methods give the same prediction regarding coexistence?

Although each of the five methods can be used to predict coexistence, the experimental approaches required for those methods are different, and it is not clear that the methods would yield the same predictions (or values of ND and RFD) if applied to the same study system. Here we use numerical simulations to investigate whether four of those methods, when implemented as shown in Figures 1, 2, 3, and 5, lead to the same prediction regarding coexistence and give the same estimates of ND and RFD. We could not include both the method based on MacArthur’s CRM and the method based on Tilman’s CRM since these mechanistic models have incompatible assumptions – the resources in MacArthur’s CRM have their own population dynamics while the resources in Tilman’s CRM are abiotic and governed by a constant rate fo supply. We chose to use numerical simulation for this demonstration since we are unaware of any empirical dataset that has been, or could be, analyzed using more than two of the methods. The numerical simulations were based on Tilman’s consumer-resource model (Tilman 1977) with two species of phytoplankton competing for two essential resources (See Supporting Information A). For each set of resource conditions, we performed numerical simulations that represent four distinct methods: 1) fitting the Lotka-Volterra model to monocultures and a co-culture (Figure 1), 2) the sensitivity method (Figure 2), 3) the method using Tilman’s CRM (Figure 4), and 4) the NFD method (Figure 5).

Figure 6 shows that under specific limiting assumptions, all four methods made the same prediction about coexistence and that these predictions matched the outcome based on the equilibrium condition from simulations. Across the different resource conditions that we explored, the two species were predicted to coexist when the resource supply conditions caused each species to be limited by a different resource, consistent with resource-ratio theory (Tilman 1977). However, this agreement among the methods was conditional on how the Lotka-Volterra and NFD methods were parameterized. The Lotka-Volterra method only matched the predictions for coexistence from the other methods when we assumed that intraspecific competition coefficients were equal to the inverse of the carrying capacity (Supporting Information Figure A1; Section 1.2). When we estimated the intraspecific coefficients directly from the monoculture time series as they approached their carrying capacity, the method produced incorrect predictions and overestimated the range of parameter space allowing for coexistence. Similarly, the NFD method only matched the predictions for coexistence from the other methods when we 1) evaluated the slope of NFD when species’ frequencies were approaching zero and 2) used both the slope and the intercept to predict the growth rate when frequency approaches zero. Unless these conditions were met, the NFD method tended to over-or under-estimate the region of resource conditions that allows for coexistence (Supporting Information Figure A1).

**Figure 6.**
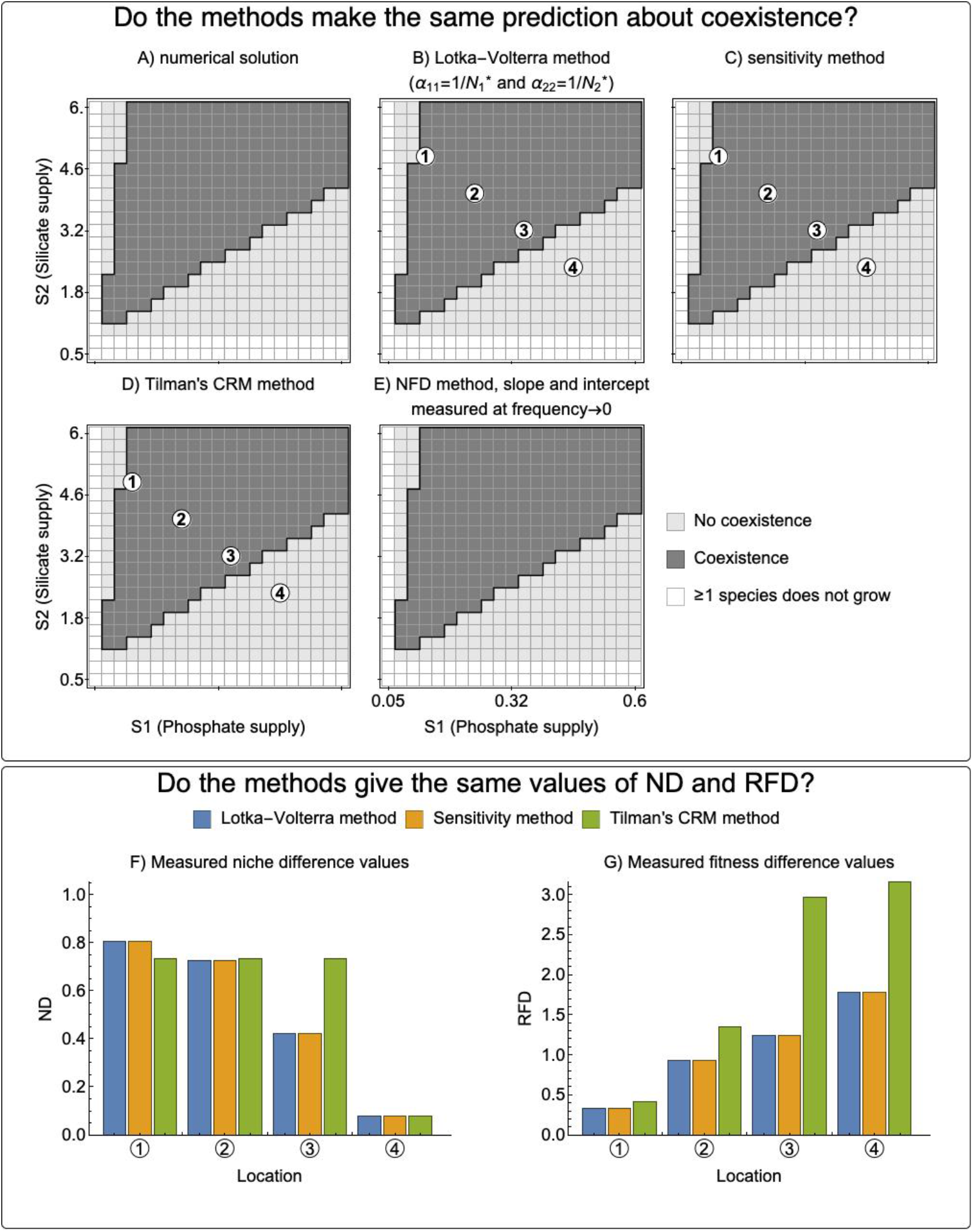
Comparison of four methods for predicting coexistence and estimating ND and RFD. The methods were compared using simulations based on Tilman’s parameterized CRM (Supporting Information A). In panels A-E, the predictions for coexistence are compared against the coexistence outcome based on numerical simulation. White shading means that at least one species does not grow under that combination of resource supply concentrations, light gray shading indicates that the method predicts that the species will not coexist, and dark shading indicate indicates that the model predicts that the species will coexist. All four of the methods give the correct predictions regarding coexistence across this region, but this is conditional on specific limiting assumptions for the Lotka-Volterra and NFD methods (see Supplement C). The methods did not give the same values for ND and RFD (f and g). The labeled locations in panels F and G correspond to marked locations in panels b-d and show that the disagreement among the methods is smaller toward the center of the parameter space that allows for coexistence. The raw RFD values from the sensitivity method were converted to the same ordering as used in the other methods (species *i* in the denominator rather than the species with the greater sensitivity). Because the NFD method cannot be used to produce values of ND and RFD that are comparable with the other four methods, only the predictions regarding coexistence are plotted.

### 1.8 Do the methods yield the same values of ND and RFD?

Although the methods gave the same predictions regarding coexistence, Figure 6 (F and G) shows that the methods do not yield the same values of ND and RFD, even when applied to the same simulated study system. The Lotka-Volterra method (using the simplification that *α_ii_*=1/*K_i_*) and the sensitivity method gave identical estimates of ND and RFD across the range of resource conditions used, but these estimates differed from the method based on Tilman’s consumer resource model. This disparity can be explained by the fact that the Lotka-Volterra and sensitivity methods assume that per capita inter- and intraspecific interaction coefficients are independent of species’ densities. Although this assumption is likely to be violated when species’ population dynamics are affected by mechanisms that produce non-linearity between population densities and growth rates, using the assumption that 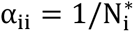 allows for accurate predictions regarding coexistence. In contrast, the method based on Tilman’s consumer resource model does not assume that interaction coefficients are independent of species densities, but instead quantifies both inter- and intraspecific interaction coefficients only at the steady-state densities predicted for monocultures that undergo invasion. This means that the interaction coefficients, and thus ND and RFD, measured according to either of the phenomenological methods (Figures 1 and 2) are unlikely to match the values predicted from a mechanistic method, even though both can correctly predict mutual invasibility.

This comparison of methods highlights an important caution, namely that estimates of ND and RFD obtained by different methods are not always comparable. Therefore, future syntheses or meta-analyses should not combine studies that measured ND and RFD by different methods. Even within a single method (e.g. the Lotka Volterra method) there can be substantial differences in the estimates of ND and RFD depending on the experimental design and how the interaction coefficients are parameterized. Similarly, estimates of ND and RFD from the mechanistic methods are dependent on the non-biological parameters used in those models (e.g. dilution rates). If a future study were to compile these values from different studies without ensuring that the same assumptions were used throughout, the results and interpretation of the synthesis would be meaningless.

## Part 2. An Empiricist’s Guide to Selecting a Method To Estimate ND and RFD

Having described and compared the foundation of each empirical method, here in Part 2 of the paper we offer practical guidance to help empiricists determine 1) which method(s) are most appropriate for their study system and 2) how much experimental effort is required for a given method. To aid our discussion, we have summarized the methods in Table 1, which is organized into three sections. The section of the table labeled ‘Decision Steps’ is a decision tree that allows an empiricist to identify the most appropriate method for their study system. The section labeled ‘Method’ directs the empiricist to the key literature needed to implement the approach. Last, the section of the table labeled ‘Experimental Requirements’ outlines key aspects of the experiments that are required to use the method.

**Table 1.**
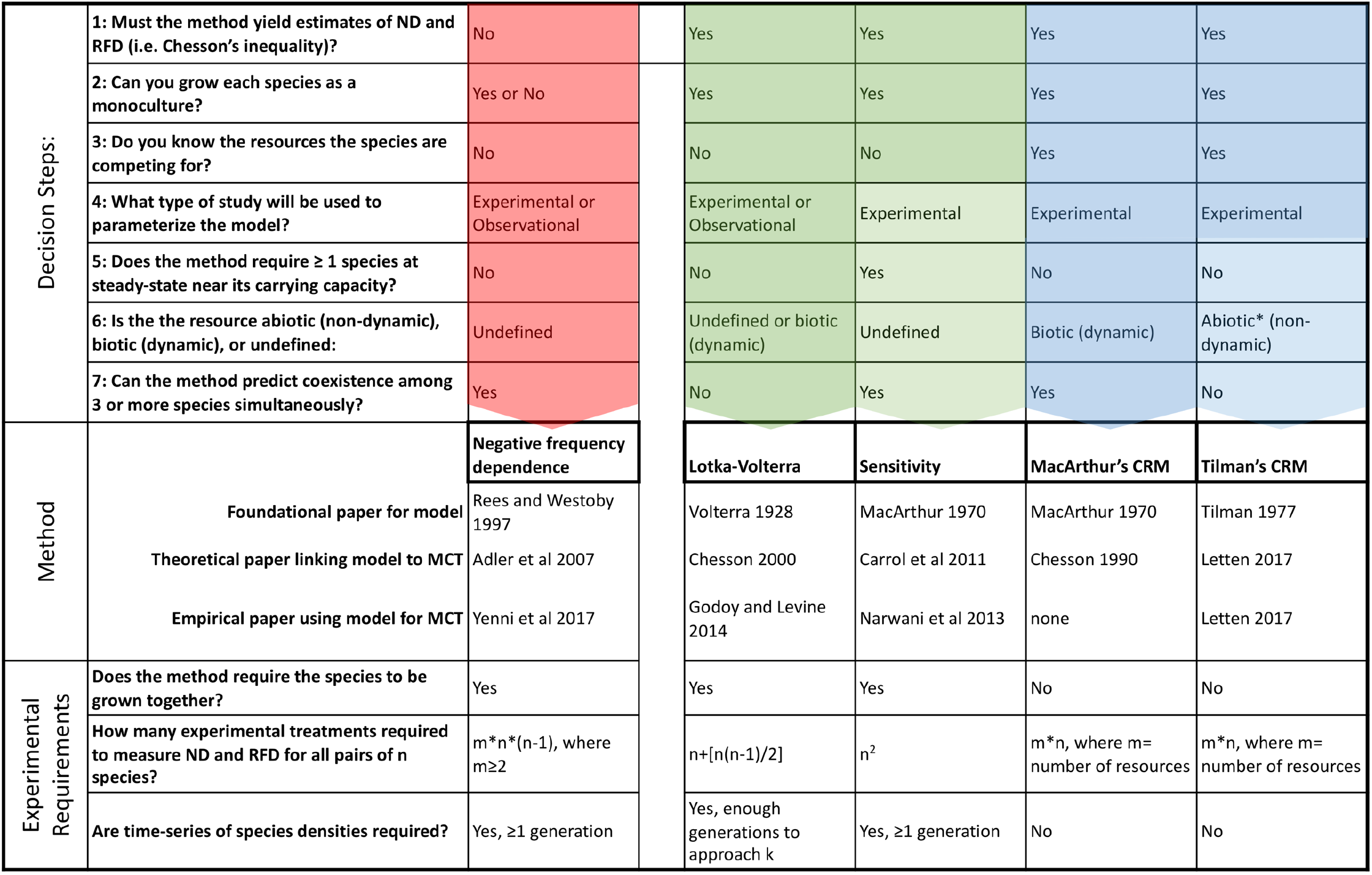
A practical guide to help empiricists determine which method(s) are most appropriate for a given study system and question. See Part 2 for a step-by-step explanation of this guide. * While consumer-resource models that include a second trophic level (e.g. predators, P*) have been developed and used empirically, these approaches have not been linked to ND and RFD.

### 2.1 Decision Steps - deciding which method to use

The first section of Table 1, ‘Decision Steps’, uses a sequence of questions about the study system to help an empiricist identify the most appropriate method for their work. The first question asks whether the method must yield estimates of ND and RFD that can be directly related back to Chesson’s inequality (Equation 1). Thus, Question 1 isolates the NFD method from all others. This distinction is important since the outputs from this method cannot be compared to the other four methods. However, the NFD method can accurately predict coexistence based on mutual invasibility and, depending on the answers to the remaining questions, it could be the most appropriate method for certain study systems. In particular, the NFD method is the only option that does not require an empiricist to grow each species alone as a monoculture (Question 2). This could be an advantage for study systems where experimental manipulations are not feasible (e.g. long-lived species, protected habitats). Several empirical studies have successfully implemented the NFD method in experiments (Levine and HilleRisLambers 2009, Chung and Rudgers 2016), and other similar studies have used NFD in observational studies (Adler et al. 2010).

The next question in the ‘Decision Steps’ is whether the empiricist knows which specific resources the species are competing for and can quantify the dependence of each species’ population dynamics on those resources (Question 3). This question separates the four methods for estimating ND and RFD into two separate groups. The phenomenological methods (Lotka-Volterra and sensitivity method) are those that are informed by directly quantifying species interactions, but which make no assumptions about the resources that species are competing for (highlighted in green). The mechanistic methods based on MacArthur’s CRM or Tilman’s CRM assume that species interact only by competing for shared resources (highlighted in blue). To use either of the mechanistic methods, an empiricist needs to know which resources define niche differences. In certain cases, it will not be possible for an empiricist to answer ‘yes’ to Question 3, because the resources required for species to grow are either not known or cannot be readily quantified (e.g. non-essential resources). When one cannot answer yes to Question 3, then the Lotka-Volterra and sensitivity methods may be appropriate because they can still quantify ND and RFD even if the empiricist does not have a good understanding of which resource(s) species are competing for, and thus, which resources define their niche. Because Question 3 is so consequential, the remaining steps are particular to either the phenomenological or mechanistic methods.

Deciding between the Lotka-Volterra method and the sensitivity method (phenomenological methods, highlighted in green), depends on the answers to whether the method must work for observational datasets (Question 4), whether it is necessary to experimentally grow each species as at steady-state near its carrying capacity (Question 5), and whether the method can be used to predict coexistence among 3 or more species simultaneously (Question 7). An empiricist working with long-lived species or in protected habitats would likely answer ‘observational’ to Question 4, eliminating the sensitivity method. In this case, the empiricist would need to decide whether it is essential to obtain values of ND and RFD compatible with the other four methods (requiring the Lotka Voltera method) or whether the NFD method could be employed to predict mutual invasibility and thus coexistence. Similarly, if an empiricist is unable to answer ‘yes’ to Question 5, she/he would be forced to use either the Lotka Volterra method or the NFD method. Question 5 could be particularly important for studies performed using slow-growing species where it is possible to parameterize the carrying capacity term from a time series of species densities, but it would take too long for the species to approach the carrying capacity to justify beginning an invasion by the other species. Lastly, the two phenomenological methods differ in terms of whether they can predict species coexistence among three or more species simultaneously (Question 7). While the Lotka-Volterra model can be parameterized to obtain all pairwise competition coefficients for a pool of species, it has not been applied to predicting coexistence of more than two species simultaneously. The sensitivity method can be used beyond pairwise species interactions (Carroll et al. 2011); however, doing so is limited to situations where all non-focal species can be considered in aggregate (e.g. species *i* invading a community of *j* + *k* + *l*).

Deciding between the MacArthur and Tilman CRM methods (mechanistic methods, highlighted in blue), is straightforward and depends on whether the resources that the species compete for are abiotic and governed by a constant rate of supply (e.g. inorganic nutrients consumed by plants) or biotic with their own population dynamics (Question 6). It is worth noting that Tilman’s R* concept has been extended to include competition mediated by predators (e.g. P*, (Tilman 1982)). However, to date, models including both predation and abiotic resource competition have not been related to Chesson’s ND and RFD. Additionally, MacArthur’s consumer resource model can theoretically work for more than two species at a time, but this has not been demonstrated for the method based on Tilman’s consumer-resource model (Question 7).

Using this decision tree, an empiricist can determine which method(s) are appropriate for their study system. Depending upon the study system or experimental constraints, an empiricist may have multiple options for which method to use. In these cases, it can be useful to consider the experimental requirements of each method (Table 1, *Experimental Requirements*) and the tradeoffs among the methods in terms of their utility as discussed in Part 3.

### 2.2 Experimental requirements

In addition to the ‘Decision Steps’ outlined in Table 1, there are important practical differences for the experimental or observational studies needed to quantify ND and RFD for each method. The most important difference in study design among these methods is whether or not they require the species to be grown together in order to make a prediction about coexistence. The NFD method and the two phenomenological methods require each pair of species to be grown together in at least one co-culture, but the mechanistic methods do not require these co-cultures. This distinction means that only the mechanistic methods can be used to make predictions about coexistence of species without performing pairwise competition experiments or analyzing time series from co-cultures. For example, consider a typical competition experiment involving a pool of three species (A, B, and C). The mechanistic methods can make predictions about species coexistence for all pairwise combinations of the species (A+B, A+C, and B+C) based solely on information about each species when grown individually. The phenomenological methods, however, require at least one co-culture for each pairwise combination of species, which means that information from monocultures and pairs A+B and A+C cannot be used to make any prediction about coexistence for the pair B+C.

The need for species to be grown together in co-culture has important implications for the total number of experimental treatments that would be required to quantify ND and RFD. Depending on the study design, experiments using the phenomenological methods can require more experimental treatments to predict pairwise coexistence among a pool of species than the mechanistic methods do. For the phenomenological methods, the number of experimental treatments required for all pairwise combinations of species increases exponentially with each additional species being considered. In contrast, for the mechanistic methods the total number of experimental treatments required increases linearly with the number of species being considered. This is because the methods based on consumer-resource models do not require any direct competition experiments in order to estimate competition coefficients (*α_ij_*), while all of the phenomenological methods require at least one co-culture for each species pair (and often more than one) in order to quantify the competition coefficients. As a result, the relative efficiency of the phenomenological versus mechanistic methods depends upon both the number of species being considered and also the number of resources. When the number of species being considered is small and the number of limiting resources is few, the difference in experimental effort can be modest. For example, to predict pairwise coexistence among a pool of four species, using the sensitivity method requires 16 experimental treatments (time series): 4 monocultures to parameterize both maximum growth rate and carrying capacity and 12 invasions to parameterize sensitivity (A invading B, B invading A, etc.). In contrast, using either of the consumer resource models (two limiting resources) would require two experiments per species for a total of 8 experiments. If the mechanistic methods require parameterizing four or more limiting resources, then the phenomenological methods may be more efficient for a pool of four species. However, for larger pools of species the difference can be substantial. Obtaining pairwise estimates of ND and RFD for a pool of 10 species requires between 55 and 180 treatments for the phenomenological methods but as few as 20 treatments for mechanistic methods.

In addition to the number of experimental treatments required for each method, it is important to consider the amount of effort and time required for each experimental treatment. Specifically, the NFD, Lotka Volterra, and sensitivity methods require time series of species densities in the experimental or observation study. In the case of the NFD and sensitivity methods, these time series may be short in duration (i.e. at least one generation) and focused only on population dynamics when species densities are very low or near the steady-state density of monocultures. However, the Lotka-Volterra method requires longer time series in order to parameterize both the interaction coefficients and carrying capacities. Longer time series in monoculture and co-culture are more easily attainable for quickly-growing species like microbes and invertebrates, but even short time series could be prohibitively arduous for slowly growing species like trees.

Ultimately, the total effort and resources required for a study is jointly determined by the method, number of species, number of limiting resources (if applicable), length of time series, level of replication, and any other design elements. Using Table 1 as a guide, an empiricist should be able to select a method and begin to design a study that satisfies their aims.

## Part 3. Tradeoffs Among Methods and Suggested Future Directions

Having explained how to select and implement the five methods, we end the paper by offering some advice for empiricists about how to navigate tradeoffs among the methods, how to compare and synthesize measurements of ND and RFD from different methods, and lastly, key future directions for implementing modern coexistence theory empirically.

### 3.1 Tradeoffs between phenomenological and mechanistic methods

Given the substantial differences in experimental design requirements and effort that are required to execute the five methods described in Part 1 of the paper, it is highly likely that empiricists will face choices that require tradeoffs when selecting a particular method for their study system. The most obvious and important tradeoffs occur between the phenomenological methods and the mechanistic methods, which differ in three important ways. First, the phenomenological methods (i.e. the NFD, Lotka-Volterra, and sensitivity methods) make no assumptions about the resources that species compete for. This could be beneficial for empiricists who can still measure ND and RFD even if they lack detailed information about the biological resources that species compete for. But the trade-off for this lack of knowledge is the need for pairwise experiments to directly quantify ND and RFD. Second, a key disadvantage of all three phenomenological methods is that they require each pair of species to be grown together in competition, which causes the total effort to increase exponentially as more species are considered. Third, the results of phenomenological experiments are specific to each pair of species tested and cannot be generalized to interactions beyond that pair. Furthermore, the predictions from the phenomenological methods are specific to the exact environmental conditions, like resource density or resource supply rates, used in that experiment and cannot be generalized outside of those same conditions.

But for those empiricists who can identify the resources that species compete for, use of the mechanistic methods allows for potentially fewer experiments that are more easily generalized to predict coexistence among all species in the focal species pool. Indeed, an empiricist who is able to answer ‘yes’ to Question 3 in Table 1 could use a mechanistic method to predict coexistence (or not) for not only the species pair of interest, but any and all species pairs of interest based solely on experiments that are performed with each species grown alone in monoculture. Importantly, the mechanistic methods also offer the ability to make predictions about species coexistence under different environmental conditions. For example, Letten et al. showed that the Tilman consumer resource model can be used to predict the ND and RFD at different nutrient supply concentrations or dilution rates (Letten et al. 2017). The ability of the mechanistic methods to handle some changes to environmental context, while limited, could be useful for predicting how anthropogenic stressors (e.g. nutrient pollution) are likely to affect species coexistence. The ability to make predictions about combinations of species without the need to perform all pairwise competition experiments has already been touted as a benefit of the mechanistic models (Tilman 1982), and it could be useful for addressing certain ecological questions that do not always lend themselves well to manipulative experiments (e.g. invasions by introduced species, coexistence of rare or endangered species).

### 3.2 Comparing and synthesizing measurements of ND and RFD

To date, only three of the four methods proposed for measuring niche and relative fitness differences have been used empirically. No one, to our knowledge has used the MacArthur consumer-resource model to quantify ND and RFD in any real system, despite publications showing that it is possible. That means that most of our inferences about ND and RFD that have been measured empirically stem from the phenomenological methods. Furthermore, we are unaware of any study that has applied more than one method to the same empirical study system. As such, we have no way to compare the performance of the methods empirically. Therefore, we believe an important avenue for future research is to focus on measuring ND and RFD using more mechanistic models, and for studies that measure ND and RFD using different methods in the same study system so that we can compare results and attempt to demonstrate equivalence or non-equivalence of these methods.

Even as we call for more mechanistic experiments and comparative studies, we caution against the inevitable urge to synthesize ND and RFD in an informal data synthesis or more formal meta-analysis. Although we have shown that all five existing methods should correctly predict the qualitative outcome of coexistence, the methods are by no means mathematically or practically equivalent. In some cases the methods will not yield the same ND and RFD, even when applied to the same species and environmental conditions. Indeed, given the differences in how the methods are implemented (Figures 1-5), there is no reason to expect, *a priori*, that the quantitative values of ND or RFD measured for a particular group of organisms using one method will produce quantitatively similar values of ND (or RFD) for that same group of organisms using a different method. As such, the methods are not directly comparable, and the measurements they produce should not be mixed-and-matched to produce some synthesized estimate of the niche or fitness difference for, say, grassland plants.

### 3.3 Future directions for implementing modern coexistence theory

In our view, there are at least two important new directions that work on species coexistence must go if Chesson’s modern coexistence theory is to become widely implemented and more practical. First, each of the empirical methods described in this review are focused on fluctuation-independent mechanisms. To also include fluctuation-dependent mechanisms of coexistence in Chesson’s framework, we need to expand the scope of the five methods reviewed here or even develop new empirical methods. To our knowledge, there have been limited empirical studies that explicitly quantify the fluctuation dependent mechanisms, i.e. relative nonlinearities and storage effects (but see (Angert et al. 2009, Letten et al. 2018)). Even so, it is well-known that environmental fluctuations mediate species coexistence in some empirical systems (Caceres 1997, Jiang and Morin 2007) and any modern theory of coexistence is incomplete without them. It is also important to note that all of the methods developed to date are only applicable to competitive communities, and cannot be applied to cases where species facilitate the growth of each other (but see (Bimler et al. 2018)).

Second, empirical studies on coexistence need to move beyond prediction of pairwise species interactions. Several authors have recently emphasized that modern coexistence theory is under-developed for multi-species systems (Carroll et al. 2011, Levine et al. 2017). In theory, the pairwise competitive hierarchy between species *i* versus *j* and *j* versus *k* might not directly translate to species *i* and *k*, particularly when these species are engaged in intransitive competition or higher-order interactions (Levine et al. 2017). In fact, none of the three phenomenological methods (the NFD, Lotka-Volterra and sensitivity methods) can deal with intransitive competition or higher-order interactions. Importantly, the emphasis to date on pairwise interactions and experimentation means that intransitive competitive interactions and higher-order interactions, if present, are unaccounted for in our understanding. Chesson’s coexistence framework has been a major advance for understanding coexistence among pairs of species, and how to expand this framework to multi-species systems should be a priority for the field.

## Author Contributions

All three authors designed the synthesis and wrote the manuscript, FHC performed the analytical derivations, CMG wrote the numerical simulation code and drafted the conceptual figures.

## Supporting Information

In the supporting information section we provide: (A) Numerical Simulation of Experiments To Measure ND and RFD and Predict Coexistence, (B) Relating the Sensitivity Method to Chesson’s Definition of ND and RFD Using the Lotka Volterra Model, and (C) Relating the Negative Frequency Dependence Method to Chesson’s ND and RFD. Annotated computer code is provided as a separate file.

## Supporting Information A: Simulation of Experiments To Measure ND and RFD and Predict Coexistence

In this supplement, we performed numerical simulations to compare the outcomes from three methods for measuring ND and RFD and also the NFD method for predicting coexistence. We used Tilman’s parameterized consumer-resource model for two species of phytoplankton competing for essential and non-substitutable resources (Tilman 1977). Annotated code for the simulations is provided in a supplemental file. Simulations were performed using the function NDSolve in Mathematica 11.2 (Wolfram Research), employing a variable step size. For each set of resource supply concentrations, we performed four simulations: (1) species 1 as a monoculture, growing from rare to near its equilibrium density; (2) species 2 as a monoculture, growing from rare to near its equilibrium density; (3) species 1 at its equilibrium density, with species 2 invading from rare; (4) and species 2 at its equilibrium density, with species 1 invading from rare. Additionally, we performed numerical simulation where both species are introduced at low densities and asked whether they coexist at the equilibrium. For each set of simulations, we manipulated the supply concentration of the two resources in order to determine whether the methods consistently agree.

We implemented the Lotka-Volterra method using information from all four simulations described above. Simulations 1 and 2 were used to estimate *r_i_*, *K_i_*, and intraspecific interaction coefficients *α_ii_*. We estimated the intraspecific interaction terms using two different approaches (Section 1.2). First, we estimated *α_ii_* as the slope of the relative growth rate versus population density (sign reversed) as the monoculture simulations approach equilibrium (Figure 1). Alternatively, we used the assumption that *α_ii_*=*1/K_i_*. We then used the parameter values from the monocultures, along with simulations 3 and 4, to solve Equation 2 when each species is at low density and the other is near equilibrium. We used all four interaction coefficients to calculate ND and RFD using Equations 3 and 4. We implemented the sensitivity method following Equations 5 through 7, using output from all four simulations. The raw RFD values from the sensitivity method were converted to the same ordering as used in the other methods (species *i* in the denominator rather than the species with the greater sensitivity).

Next, we compared the four methods including the sensitivity method, the method based on Tilman’s CRM, the Lokta-Volterra method, and the NFD method using the numerical simulations described above. Under specific assumptions, the methods gave the same prediction regarding coexistence (Figure A2), though the methods did not produce consistent estimates of ND and RFD (Figure 6).

As described in Appendix C, the NFD method cannot be used to get ND and RFD estimates that are consistent with the other methods, but nonetheless this method can be used to predict coexistence based on the same criterion. However, as shown in Figure 5 (using the Lotka-Volterra model), accuracy of the NFD method depends on the range of frequencies used to get the slope and elevation. To illustrate how the non-constant NFD slope is problematic in predicting species coexistence, we used the simulations of mutual invasion (simulations 3 and 4), described above, to construct pairs of each species’ frequency and their growth rate in a saturated community. For all of these simulations, we used only supply concentrations of the resources that are known to allow for coexistence. For each value of a species frequency between 0 and 1, we calculated the slope of growth rate versus frequency. Figure A1 shows that this slope is not constant and actually changes sign depending on the species’ frequencies used. Thus, using only the slope of the NFD relationship is inadequate to predict coexistence.

Next, we used both the slope and elevation from the NFD method to extrapolate to frequency of 0 and predict whether the species is capable of invasion from rare (Figure A2 panels A and B). Figure A2 shows that for supply conditions known to allow coexistence, the accuracy of the predictions from the NFD method depends on the range of frequencies over which the slope of NFD was measured. We discuss two points (A and B in Figure A2) to explain this effect. At the point Labeled “A”, the slope of NFD for species 1 predicts a positive growth rate as frequency approaches 0, but at the complementary frequency of species 2, the slope of species 2’s NFD predicts a negative growth rate when rare. However, based on the other 3 methods, numerical simulation, and Tilman’s resource ratio theory, the species are predicted to coexist. Thus, measuring NFD under the red regions in Figure A2 will incorrectly predict exclusion even though the species will coexist. At the point labeled “B” in Figure A2, the slope and elevation of NFD for both species predicts a positive growth rate when rare. This region, depicted in blue, includes the equilibrium frequency for the two species. If an empiricist made their measurements between frequency of ∼0.05 to ∼0.85 for species 1, and used the slope of NFD, they would incorrectly predict that the species will not coexist. Since the frequency at which the species reach equilibrium depends on the resource supply ratio, there is no single frequency of the species that consistently leads to the correct predictions (Figure A2). While certain intermediate frequencies of the two species can be used to make accurate predictions, an empiricist would not know these frequencies without performing the competition experiments or examining frequency dependence across the entire range of frequencies. As a result, the only reliable way of implementing the NFD is to measure the slope and elevation for each species where its frequency approaches zero.

For the NFD method, accurate predictions required that the slope of NFD was evaluated approaching frequency of zero for each species (i.e. invasion conditions). In Figure S2 d-f, we show that evaluating the NFD slope at other frequencies leads to the wrong predictions. We used the NFD plot to evaluate coexistence at three frequencies, including near 0% (panel e and h of Figure S2), 50% (panel f and i of Figure C2) and near 100% (panel g and j of Figure C2), and either with (panel e-g of Figure C2) or without considering the elevation in addition to the slope (panel h-j of Figure C2). We see that using the NFD slope evaluated at near 0% frequency will consistently yield accurate predictions of species coexistence that match the those of the other methods.

**Supporting Information Figure A1.**
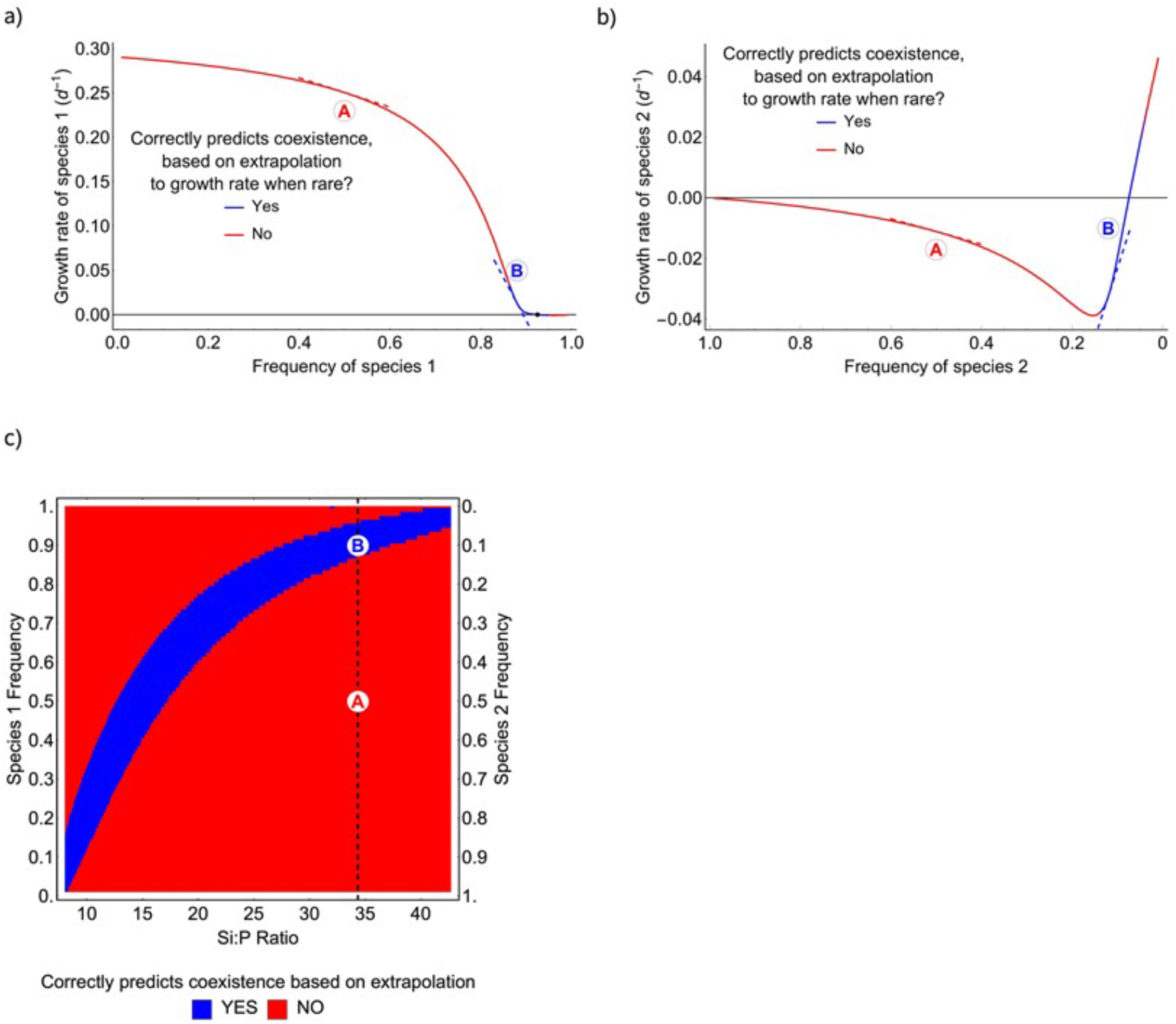
Results of simulation experiments using the NFD method. Panels a and b show per capita growth rate versus the frequency of species 1 and 2. At any frequency of the two species, the NFD method requires that we use the slope to extrapolate and estimate the growth rate when frequency approaches zero (the extrapolated vertical intercept). For frequencies where this method predicts mutual invasibility for both species, i.e. species can coexist, the lines are blue. For frequencies of the two species where the method leads to the incorrect prediction, the lines are red. Both species have positive growth rates when their frequency approaches zero, indicating that they are mutually invasible. Panel c shows the accuracy of the NFD method as a function of the supply Si:P ratio and the frequency of the two species at which the method was applied. The vertical dashed line represents the slice depicted in panels a and b. For all of the Si:P ratios shown in panel c, the species are mutually invasible and will coexist. This plot indicates that using NFD will often predict that the species will not coexist, when in fact they do coexist. This is important because without examining the full range of species frequencies in an experiment, one would not know whether and where the relationship between frequency and growth rate is non-linear.

**Supporting Information Figure A2.**
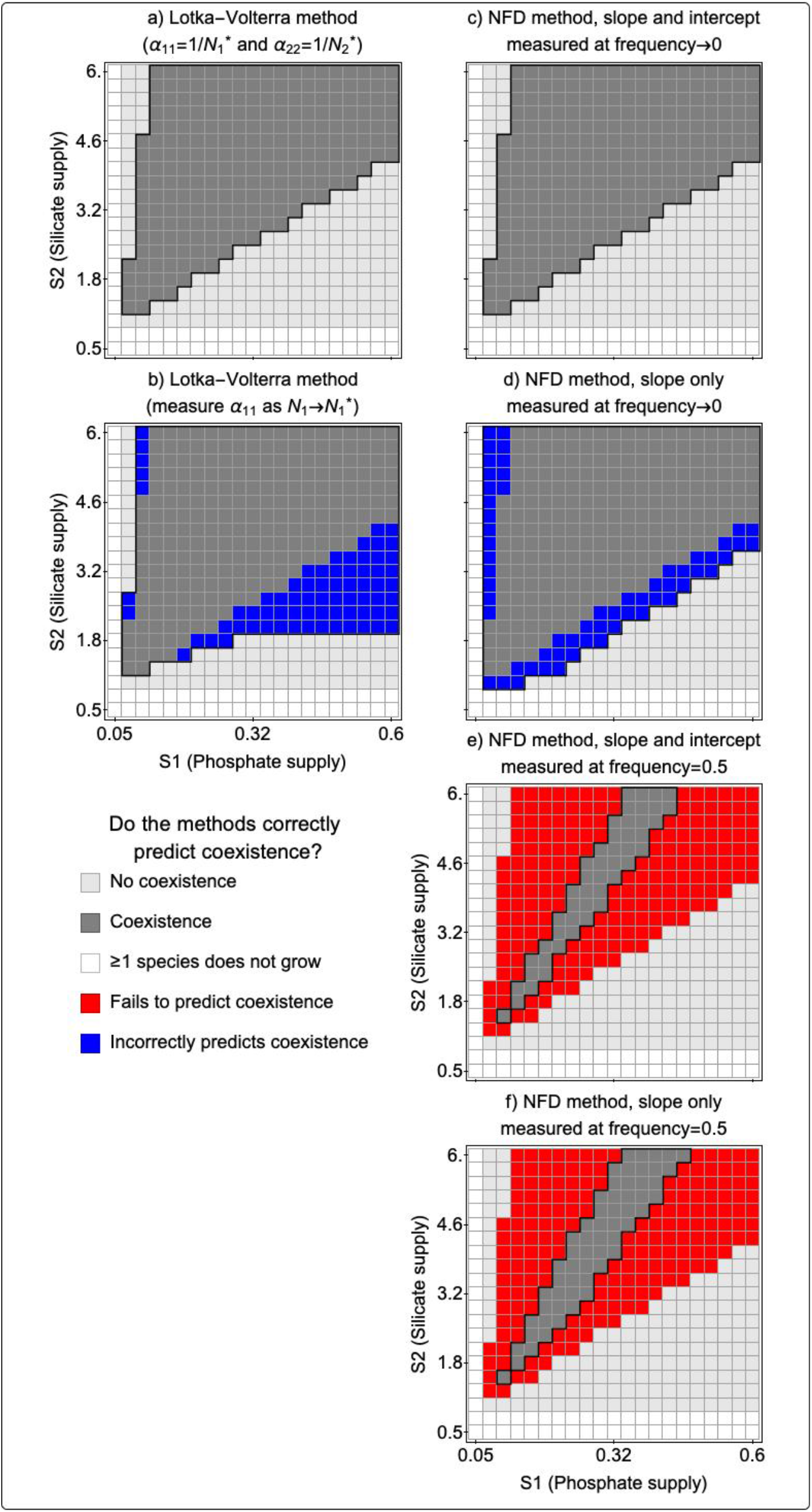
Results of simulation experiments comparing predictions from the Lotka-Volterra and NFD methods.

## Supporting Information B: Relating the Sensitivity Method to Chesson’s Definition of ND and RFD Using the Lotka Volterra Model

Here we show that sensitivity method is identical to the Lotka Volterra method given the specific limiting assumptions of the sensitivity method. To do this, we derive the sensitivity metric (*S_i_*) from the Lotka-Volterra competition model (Equation 2). The **μ*_i_* in Equation 5 is the maximum per capita growth rate in monoculture, equal to *r_i_*in Equation 2. The **μ*_ij_* is the invasion growth rate, so that we can replace *N_j_* with species j’s carrying capacity, *K_j_*, and replace *N_i_* with 0, so that **μ*_ij_*=*r_i_*(1-*α_ij_ K_j_*). Using this substitution, we show in Equation B1 that the sensitivity metric (*S_i_*) is the equilibrium density of species *j* (*K_j_*) multiplied by the *per capita* competition coefficient (*α_ij_*).

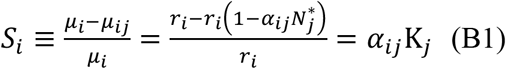

Since the intraspecific competition coefficients in the Lotka Volterra model are equal to the inverse of the equilibrium population density for the monoculture 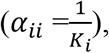 the sensitivity metric can be shown to be equivalent to the ratio of interspecific to intraspecific interaction coefficients (Equation B2).

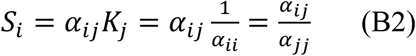

From this substitution, we can relate the sensitivity metric to Chesson’s ND (Equation B3), RFD (Equation B4), and use these estimates to assess the conditions for coexistence (Equation

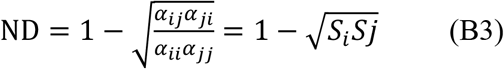

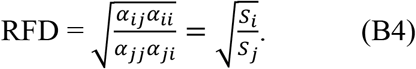

## Supporting Information C: Relating the Negative Frequency Dependence Method to Chesson’s ND and RFD

Here we show that in order for the NFD slope to be constant, the community density must be both saturated and fixed across all frequencies of the species. To do so, we attempt to derive the NFD slope and intercept from the two species Lotka-Volterra competition model (Equation 1). Since there is no variable representing each species’ frequency in the Lotka-Volterra model, we have to assume a fixed community density, *B*. This assumption also satisfies the assumption of the NFD method that the community density is always saturated. Fixing the community density makes the interspecific density dependence, *α_ij_*, equivalent to frequency dependence (Adler et al. 2007), and allows species’ frequency to be represented by *N_i_*/*B*. The two-species Lotka-Volterra competition model can then be rewritten as follows

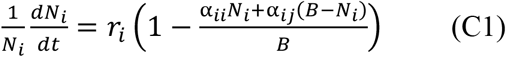

where *B* is the fixed community density and one unit decrease of *N_i_* will lead to one unit increase of *N_j_*. From Equation C1, we derive the NFD slope and intercept in the following equations.

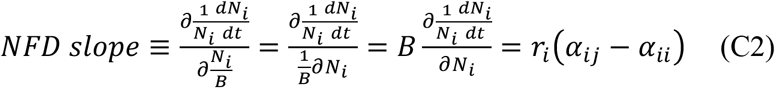

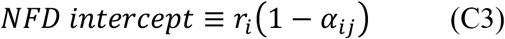

In Equation C2 the NFD slope becomes constant, which means that species’ per capita growth rate linearly depends on its frequency.

In addition, from Equations C2 and C3, we argue that both NFD intercept and slope should be used with caution in evaluating Chesson’s inequality. First, the NFD intercept represents whether species can successfully invade a steady-state population of its competitor at its carrying capacity, so it can be used to accurately assess mutual invasibility. However, neither the difference nor the ratio of two species’ NFD intercepts (Equations C2 and C3) take an analogous form to Chesson’s definition of ND and RFD. It is worth noting, however, that the slope of NFD relationship has been used to represent ND for annual plant communities (Yenni et al. 2012, Yenni et al. 2017). Thus, while the NFD method can correctly predict mutual invasibility, the NFD intercept and slope should not be interpreted as RFD and ND in order to evaluate Chesson’s inequality.

The utility of the NFD method depends on its ability to correctly predict whether species have positive growth rates when their frequencies approach zero. If the relationship between a species frequency and its growth rate is non-linear, however, then the accuracy of the NFD method is critically dependent on the range of species frequencies used by an empiricist. In Figure C1 we show that the NFD relationship is non-linear even when the underlying population dynamics are governed by the Lotka-Volterra model. The result of this non-linearity is that, depending on the range of species’ frequencies used to estimate the NFD slope and intercept, this method can give inaccurate predictions.

**Figure C1:**
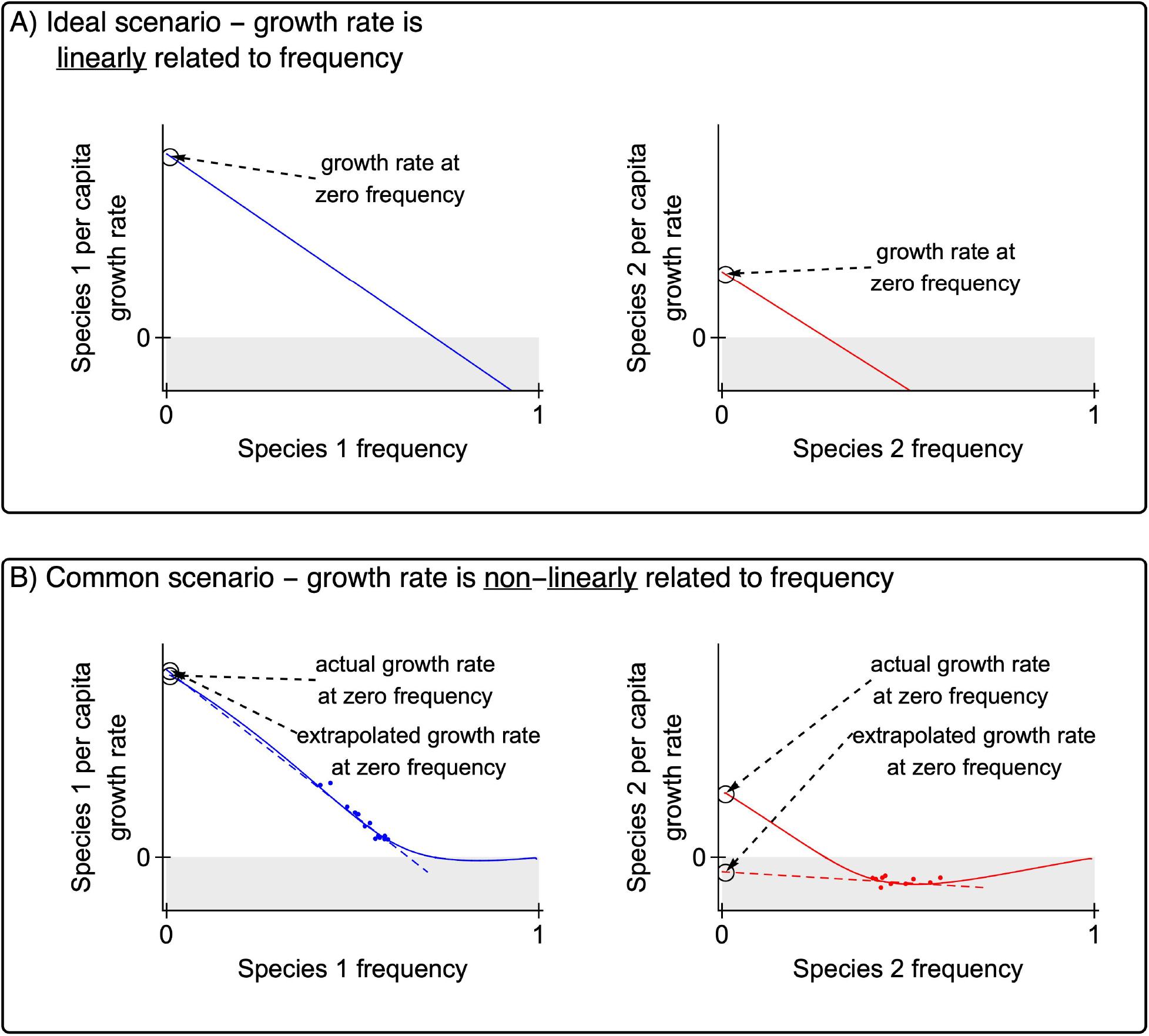
Panel A shows a hypothetical situation where species frequencies are linearly related to their growth rate and an empiricist can extrapolate to predict growth rates at frequency of zero and diagnose mutual invasibility. Panel B shows the more likely scenario in which growth rates are non-linearly dependent upon species frequencies. These plots were made using numerical simulation of the two-species Lotka Volterra Model, using parameter values that should allow for coexistence (at frequency of 0.72 for species 1). The points in Panel B represent empirical measurements collected at intermediate frequency of both species. Using those measurements and extrapolating to zero frequency yields the incorrect prediction that the species will not coexist.

